# A Radiomic Approach to Clinical MRI Refines the Thalamus-Cognition Link in Multiple Sclerosis

**DOI:** 10.1101/2025.06.05.657698

**Authors:** Korhan Buyukturkoglu, Lin Lu, Levi Davis, Renan E. Orellana, Charles C. White, Rongyi Sun, Sinem Ozcelik, Nina M. Isenstein, Kaho B. Onomichi, Rifat Iqbal, Binsheng Zhao, Yaakov Stern, Burcu Zeydan, Orhun H. Kantarci, Claire S. Riley, Philip L. De Jager

**Affiliations:** Columbia University; MSKCC; Mayo Clinic

**Author notes:** Corresponding author: Korhan Buyukturkoglu The Neurological Institute-Columbia University Irving Medical Center 710 W 168th St. NI-09-900F Room 102 New York, New York 10033. **Funding:** The work presented here is supported by National MS Society Harry Weaver Award to KB-JF-2208-40306, NMSS grant SI-1903-33766 to PLD and NIH/NIA grant to YS 5R01AG038465.

**Keywords:** Radiomics, thalamus, thalamic nuclei, cognition, SDMT, PASAT, DSST

## Abstract

**Background and Objectives:** Radiomics extracts imaging features that may not be detectable through conventional volumetric analyses. Given their role in multiple sclerosis (MS), we applied radiomics to thalamic nuclei and examined their associations with cognitive performance.

**Methods:** A total of 601 individuals were included (342 people with MS-PwMS from two cohorts, and 259 healthy controls-HC). Radiomic features (RF) and volumes were extracted from the whole thalamus, five thalamic nuclei, and the putamen segmented on 3D T1-weighted images. Cognitive performance was assessed using the Symbol Digit Modalities Test (SDMT) and Paced Auditory Serial Addition Test (PASAT) in PwMS, and the Digit Symbol Substitution Test (DSST) in HC. In the first MS cohort, multivariate linear regression in a discovery set (N=103) identified thalamus-derived RF associated with SDMT, which were retested in a replication set (N=63). Their associations with PASAT in a second MS cohort (N=176) and DSST in HC were also evaluated. We then tested whether the same RF, when extracted from the putamen, were associated with SDMT. LASSO models assessed the combined predictive value of RF and volumes.

Figure 2 presents an overview of the involvement of different cohorts in the study, outlining the specific study objectives and the statistical analysis approaches employed.

**Results:** Twenty-eight RF–ROI pairs were associated with SDMT in the replication set (FDR<0.05). Of these, 24 were also associated with PASAT (FDR≤0.03), and 2 with DSST. Only ventral nuclei volume showed replicated associations among volumetrics. Only 4 putamen-derived pairs were associated with SDMT (FDR=0.04). LASSO results confirmed RF outperformed volumes.

**Discussion:** RF extracted from the thalamus are strongly associated with cognitive performance in PwMS, outperforming volumetric measures and supporting their potential as sensitive imaging biomarkers.

## Introduction

Cognitive impairment is a debilitating symptom of multiple sclerosis (MS) (1). Magnetic resonance imaging (MRI) offers valuable insights into MS-related brain changes; however, despite recent advances, neuroimaging signatures that fully explain the causes and progression of cognitive impairment remain insufficient. One contributing factor may be the challenge of capturing the heterogeneity of MS using conventional/structural imaging techniques. Advanced MRI sequences can reveal subtle changes and overcome this challenge, but their limited adoption in standard protocols restricts broader use. This gap highlights the need for novel imaging approaches or advanced analytical strategies that leverage widely available clinical-grade imaging data (e.g., neuroimaging data from routine visits) to extract more nuanced information, detect subtle brain abnormalities, and ultimately help development of reliable biomarkers for cognitive impairment in MS.

The thalamus is a central relay and integration hub in the mammalian forebrain (2) composed of anatomically and functionally distinct nuclei and each region contributes uniquely to brain functions including cognition (3). Among other brain regions, the thalamus has emerged as a selectively vulnerable and critically important structure in MS (4). Advanced MRI techniques such as diffusion weighted imaging (5), MRI spectroscopy (6), Positron Emission Tomography (7) and functional MRI (8) have revealed a range of abnormalities in the thalamus associated with cognitive impairment in MS. While these techniques provide valuable insights, they are often expensive, technically demanding, not widely accessible, and in some cases, slightly invasive (e.g., PET). As a result, their application in large-scale or routine clinical settings is limited.

On the other hand, conventional MRI sequences (such as T1 and T2 weighted imaging) provide a wealth of data as they are typically acquired during routine/clinical visits. These images can reveal thalamic atrophy and lesions, both of which are associated with cognitive deficits in MS (9,10). However, they may not fully capture the subtler pathological changes (such as alterations in tissue organization, cellular density, and microstructural integrity) which may be also linked to cognitive impairment. Furthermore, conventional markers often reflect damage that is already well advanced. As such, they may be insufficient as predictive biomarkers of future cognitive decline.

Radiomics, a quantitative image analysis approach (11) enables the extraction of high-dimensional features from medical images. Compared to advanced image analysis modalities, it is more scalable and accessible, yet capable of detecting morphological and microstructural alterations (12). Techniques such as texture analysis quantify tissue variations that may precede volume loss (13), while features measuring intensity distribution, spatial complexity, and tissue heterogeneity can reveal microstructural disorganization and local signal variability potentially reflecting pathological processes like inflammation or demyelination not visible on conventional imaging (14).

Radiomics has shown significant potential in oncology, where they predict treatment response and disease progression earlier than volume-based measurements (15–18). This approach has also contributed to understanding neurologic conditions like Parkinson’s (19) and Alzheimer’s disease (20) by identifying novel radiological markers and providing insights into disease mechanisms.

Despite its successes in other fields, its application in MS remains limited. Most MS-related radiomics studies focus on lesions or classification problems, such as distinguishing MS from similar diseases (21–25) (Please see Inglese et al for a comprehensive review) (26). Only a few studies have investigated the cognitive aspects of MS so far (27). One notable study integrated clinical data, multi-sequence MRI based radiomics, and structural features to identify cognitive performance and predict its progression in people with MS (PwMS), using the Symbol Digit Modalities Test (SDMT) as the cognitive outcome. Models that integrate clinical data, multi-sequence lesion radiomics, and structural features yielded excellent results in identifying cognitive impairment and predicting cognitive worsening (28).

Motivated by these insights, in this study we aimed to investigate whether radiomics can uncover novel, clinically relevant imaging signatures linking cognitive performance to thalamic nuclei, going beyond conventional volumetric analyses and adding value to routinely acquired clinical MRI scans. Since information processing speed, attention, and working memory are among the most commonly affected cognitive domains in MS (29) we selected the SDMT as our primary cognitive outcome, as it is the most widely used screening tool for evaluating performance in these domains (30).

To achieve our aims, we first analyzed data from 166 PwMS enrolled in an ongoing longitudinal study (MS Snapshot) (31). Using multivariate linear regression analysis, we identified radiomic features related to SDMT scores in a subset of this cohort (discovery set, N = 103). We then tested the replicability of the features in a replication set consisting of participants from the same cohort who were recruited later into the study (N = 63).

To test the anatomical specificity of our results, we used the putamen (another deep gray matter structure of comparable size to the thalamus) as a control region. As the putamen is predominantly associated with motor functions (32), we hypothesized that the thalamus derived radiomic features associated with SDMT performance would not demonstrate the same associations when extracted from the putamen. Furthermore, we aimed to assess whether radiomic features capture shared neural substrates underlying performance within the same cognitive domain when evaluated using different tests. Specifically, we examined whether key radiomic features identified in our initial discovery-replication analyses as associated with SDMT performance were also associated with performance on the Paced Auditory Serial Addition Test (PASAT) (33). Although the two tests engage partially distinct cognitive processes potentially driven by different brain regions (with PASAT additionally requiring auditory processing and calculation skills) they both tap into overlapping cognitive domains (information processing speed, working memory, and attention). Therefore, we hypothesized that thalamus-derived radiomic features would reflect similar abnormalities linked to both. This analysis was performed using an independent cohort of 176 PwMS from the Mayo Clinic (34).

Lastly, to assess disease specificity, we analyzed healthy controls (HC) from the Reference Ability Neural Network (RANN) Study (N = 269) (35) who completed the Digit Symbol Substitution Test (DSST) (36), a cognitive measure analogous to the SDMT. Here we applied Least Absolute Shrinkage and Selection Operator (LASSO) regression with cross-validation to both the MS Snapshot cohort and the HC to test specificity and to determine whether combining radiomic and volumetric features enhances prediction accuracy.

Overall, in this study we aimed explore the potential of radiomics to improve the sensitivity of measures derived from T1-weighted/clinical MRIs beyond what volume measurements provide. We selected the thalamus and its nuclei as our target regions because thalamic abnormalities are commonly observed in MS and highly correlated with disease symptoms.

## Materials and Methods

### Description of the Participants and Data

The institutional review board of Columbia University Irving Medical Center (CUIMC) and the Mayo Clinic approved the study protocols, and all participants provided written informed consent. The demographic and cognitive characteristics of the cohorts is presented in Supplementary Table 1.

*MS Snapshot Cohort:* We accessed MRI data collected as part of routine clinical care from PwMS involved in an ongoing longitudinal study (MS Snapshot) (31) which includes prospective collection of the brain and spinal cord MRIs. A total of 166 PwMS (mean age 48.5±13, 115 Female) who had both cognitive testing and clinical MRI collected within maximum 6 months of each other were selected. On average, the time gap between the MRI acquisition and the cognitive tests was 76 days (max=180 days, min=1 days).

Cognitive function was evaluated remotely via secure Zoom link during the participants’ clinical telehealth visits between 2020 and 2024 (31). The SDMT (37) is generally considered to be the most sensitive cognitive measure for use in MS and is the most-utilized cognitive measure in MS clinical trials (30,38). It assesses cognitive domains including processing speed, attention, working memory, and visual scanning by having individuals match symbols to corresponding numbers using a reference key. It has been shown that remote and in-person administration of the SDMT yield comparable results (31,39).

The majority of the MR images (N=113) were obtained using a 3 Tesla GE Discovery MRI scanner (GE Healthcare, Wauwatosa, WI). Additionally, data from other MRI scanners from other centers were included.

*Mayo Clinic Cohort:* In this cohort participants were prospectively enrolled from an outpatient MS clinic (34). A total of 176 PwMS (mean age 48±13 years, 127 Female) who visited the clinic between 2021 and 2024 were included. Eligible brain MRIs were selected from scans acquired as part of routine clinical care.

For this data set, the PASAT (40) as a component of the Multiple Sclerosis Functional Composite (MSFC) has been used to assess cognitive function. The PASAT measures attention, auditory information processing speed, concentration, working memory, cognitive flexibility, and calculation skills which are partially regulated by the thalamus, by having participants listen to a series of single-digit numbers and add each new number to the one immediately preceding it.

The majority of the MR images (N=111) were obtained using a 3 Tesla Siemens MRI scanner (Siemens Healthineers, Germany). Additionally, data from other MRI scanners were included. The median time between cognitive testing and brain MRI was 3 days (interquartile range=0-57 days).

*RANN Study Cohort:* We processed an additional set of MRI data collected from 259 non-MS/healthy participants (mean age 52±16, 146 Female) in the RANN study (35). All RANN participants were screened for serious psychiatric or medical conditions, poor hearing and vision, and any other impediments that could hinder MRI acquisition. In addition, older participants were screened for dementia and mild cognitive impairment using the dementia rating scale.

Among the twelve cognitive tests were administered as part of the RANN study, the Digit Symbol Substitution Test (DSST) (36) was selected for this study due to its similarity to the SDMT. DSST, entails converting digits into symbols (while SDMT requires translating symbols into digits). Both tests aim to assess cognitive processing speed and attention, employing slightly distinct methods to achieve this goal.

Image acquisition was performed using a 3T Philips Achieva Magnet (Philips Healthcare, Best, The Netherlands).

In each cohort, MRI protocols included various structural and functional sequences; however, only the 3D T1-weighted images (all ≤ 1mm, isotropic) were used for volumetric and radiomic analyses in this study, as these images were consistently available across all cohorts. A detailed list of scanners and acquisition parameters used in this study can be found in the Supplementary Material.

### MRI Data Analysis

The Fluid-Attenuated Inversion Recovery (FLAIR) and 3D T1-weighted images were reviewed by two neurologists (SK and RS) and a neuroscientist (KB) to assess thalamic lesions. The presence of a lesion within otherwise healthy brain tissue can significantly impact radiomic features by altering the intensity, texture, and shape characteristics of the region of interest (ROI). Therefore, patients with thalamic lesions were not included in this study.

As a first step of MRI analysis the N4 bias correction algorithm was utilized to eliminate non-uniform low-frequency intensity in the images. Considering the potential effects of global T1 hypointense lesions on thalamus segmentation (41,42) we used lesion-in painted images for generating the ROI masks. Automated lesion segmentation was performed using SAMSEG (43) a module within FreeSurfer followed by lesion inpainting performed using FSL (FMRIB Software Library). Following that, the thalamic nuclei were segmented using FreeSurfer thalamic subnuclei segmentation tool (44). Right and left individual thalamic nuclei masks were merged to create ventral (combinations of ventral anterior, ventral anterior magnocellular, ventral lateral anterior, ventral lateral posterior, ventral posterolateral, ventromedial nuclei) medial (combination of paratenial, medial ventral, mediodorsal medial magnocellular, mediodorsal lateral parvocellular nuclei), lateral (laterodorsal, lateral posterior), intralaminar (central medial, central lateral, paracentral, centromedian, parafascicular) and pulvinar (combination of pulvinar anterior, pulvinar medial, pulvinar lateral, pulvinar inferior) nuclei masks (see Figure 1). We used the left and right whole thalamus masks generated by the same tool instead of those provided by FreeSurfer’s standard recon-all stream, as the former have been reported to yield more precise thalamic segmentation results (44).

**Figure 1.**
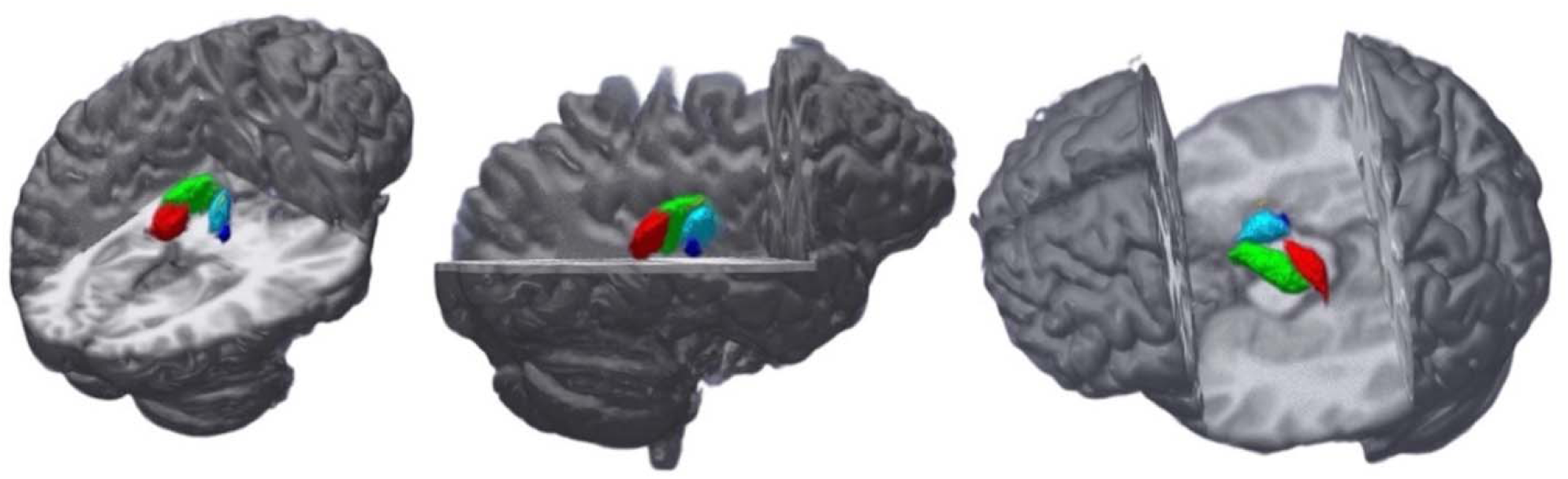
The thalamic nuclei selected as region of interests in this study. Masks of region of interests (ROIs) of the study are highlighted in different colors. Red: Left Pulvinar, Green: Left Ventral, Yellow: Right Intralaminar, Dark Blue: Right Lateral, Light Blue: Right Medial Nucleus.

**Figure 2.**
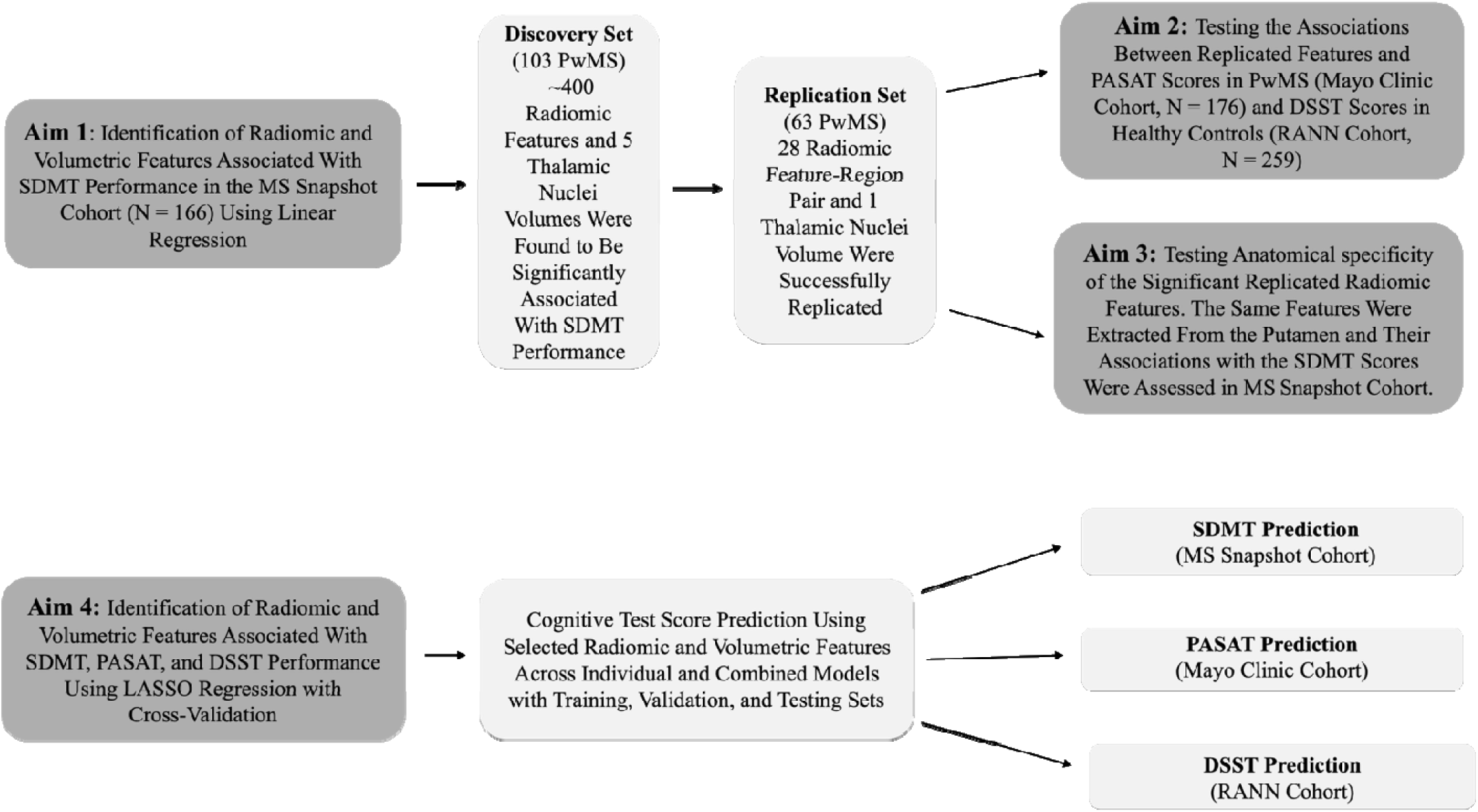
Overview of the involvement of different cohorts in the study, specific study objectives and the statistical analysis approaches employed. Abbreviations: SDMT: Symbol Digit Modalities Test, PASAT: Paced Auditory Serial Addition Test, DSST: Digit Symbol Substitution Test, LASSO: Least Absolute Shrinkage and Selection Operator.

Total intracranial brain volume (ICV) was measured using FreeSurfer version 7.3.2.50. To account for individual head size differences, volumes were normalized by ICV for each participant prior to statistical analysis.

### Radiomic Feature Extraction

Radiomic features including first order statistics (19 features), 26 3D and 2D shape-based features, 24 gray level co-occurrence matrix, 16 gray level run length matrix, 16 gray level size zone matrix, 5 neighboring gray tone difference matrix and 14 gray level dependence matrix features were computed for 6 ROIs (bilateral whole thalamus, pulvinar, medial, ventral, lateral and intralaminar nuclei) using PyRadiomics package (18). All feature classes, with the exception of shape can be calculated on either the original image and/or a derived image, obtained by applying one of several filters such as wavelet or Laplacian of Gaussian filters-LOG filters. With the use of these filters the total radiomic feature number per subject/per ROI went up to 1734. For each ROI mask radiomics features were firstly extracted from both the left and right and then averaged. We then applied ComBat harmonization (NeuroCombat Python Package, V 0.2.12) to remove batch effects due to differing locations and type of MRI scanners (45). Following this step, each feature value was standardized using z-score normalization.

## Statistical Analyses

### Identifying Radiomic and Volumetric Features Associated with SDMT Scores

Our first aim in this study was to assess whether thalamus derived radiomic or volumetric features extracted from clinical MRIs are related with SDMT performance in PwMS. To evaluate the robustness of the associations, we applied a split-sample validation strategy using discovery, replication, and meta-analysis phases. The discovery set included 103 PwMS enrolled early in the study, while later-enrolled participants (N=63) comprised the replication set.

To prepare the data, we reduced dimensionality and addressed multicollinearity by retaining only the first feature from each group of highly correlated radiomic features (r > 0.90), resulting in approximately 400 radiomic features per ROI. Each feature was analyzed independently in the discovery set using linear regression models adjusted for age and sex. Features that survived false discovery rate correction (FDR < 0 .05) were tested in the replication cohort using the same model. Results from both sets were then combined via inverse-variance meta-analysis.

Our second aim was to test the anatomical specificity of the significant radiomic feature-SDMT associations. We tested this by constructing age- and sex-adjusted linear regression models that used putamen-derived radiomic features in 166 MS Snapshot participants.

### Testing the Relationships Among SDMT-Associated Radiomic Features, PASAT and DSST Scores

The third aim was to evaluate the relationship between radiomic features associated with SDMT performance in the MS Snapshot cohort and PASAT scores in an independent sample of PwMS (Mayo Clinic Sample, N=176) as well as DSST scores in HC (RANN Study, N=259). With that aim, we constructed linear regression models adjusted for age and sex using the previously identified SDMT-associated radiomic features in the MS Snapshot Cohort. The PASAT was selected as a proxy measure to assess the potential generalizability of radiomic features, specifically, whether they capture core thalamic signatures relevant to processing speed across different cognitive tasks. DSST was selected due to its similarity to the SDMT. Notably, SDMT scores were not available either in the Mayo or in the RANN datasets.

Here we also examined whether individual thalamic nuclei volumes were associated with PASAT and DSST scores.

### Integrating Radiomic and Volumetric Features to Predict Cognitive Performance in MS and Healthy Controls (LASSO analysis)

In a separate set of analyses, we applied the Least Absolute Shrinkage and Selection Operator (LASSO) regression algorithm with the aim of evaluating the combined predictive value of radiomic and volumetric features for SDMT performance in PwMS (MS Snapshot cohort). We applied the same approach to healthy controls from the RANN cohort, using DSST scores as the cognitive outcome, to evaluate the specificity of these associations to MS.

Unlike our earlier discovery-replication analyses that examined feature types independently, this approach allowed us to model joint effects and also leverage LASSO’s feature selection capabilities while controlling for overfitting through regularization. LASSO regularizes the regression model by incorporating an L1 penalty term proportional to the absolute values of the model’s coefficients. Therefore, it reduces dimensionality and helps prevent overfitting (46).

To estimate the variance explained through LASSO regression, we used a training set/testing set split approach in each cohort separately. We first randomly assigned 70% of our subjects into a training data set and the remaining 30% into a testing dataset. We then performed LASSO analysis on the training set to develop our predictive model, which we subsequently tested on the testing dataset. Note that, this 70%–30% training–testing split is independent of the earlier discovery (N=103) and replication (N=63) cohort split.

To assess the predictive value of the derived model, we used it to generate predicted cognitive test scores based on the radiomic and volumetric features and then compared these predictions with the actual cognitive test performance in the testing dataset. To quantify the predictive value of the model, we calculated a correlation coefficient and r^2^ between the predicted and observed values using a Pearson correlation test. Given that the predictive performance of the derived model is sensitive to the random dataset split, we iterated this process 1000 times to calculate a median r^2^ value and median p-value for the correlation between the predicted and observed score.

To compare the relative r^2^ of different variable sets, we ran the simulations on a series of models for each ROI separately, including: 1) models consisting of solely radiomic features, 2) models using only individual thalamic nuclei volumes, and 3) models combining radiomic and volumetric features for each ROI. Details of the LASSO analysis are provided in the Supplementary Material.

Note that to predict cognitive test scores using the volumes of individual thalamic nuclei (such as exploring the association between pulvinar nucleus volume and SDMT), we applied linear regression instead of LASSO regression due to the small number of variables.

## Results

### Description of participants

The MS Snapshot cohort (N=166) is a longitudinal study of PwMS with prospective brain autopsy; these participants were divided into a Discovery and a Replication sample set. We also used an external set of PwMS from the Mayo Clinic (N=176) for generalization of our results as well as participants in the longitudinal study of cognitive reserve, RANN (N=259) (see Supplementary Table 1 demographic and clinical characteristics of all cohorts).

### Radiomic Features and Thalamic Volumes Related to SDMT in PwMS

Following preprocessing and our dimensionality reduction step (see Methods), approximately 400 radiomic features extracted from the thalamus and its nuclei were evaluated in our discovery analysis based on 103 participants from the MS Snapshot study with MRI data from clinical scans as of March 2022. The 63 participants who entered the study after this date and before June 2024 were used in the replication analysis. These features were derived from various image transformations (e.g., original, wavelet, logarithmic, gradient) and categorized into shape, first, second, higher order, and texture-based features. Morphological features characterize the shape and size of the ROI, including parameters like effective diameter, surface area, and volume. First-order features describe the statistical distribution of voxel intensities using measures such as maximum, minimum, standard deviation, skewness, and entropy. Texture features encompassing second- and higher-order statistics capture spatial relationships within the image. Second-order features assess local intensity patterns (e.g., energy, entropy, contrast, uniformity), while higher-order features describe more complex spatial structures such as heterogeneity and roughness. These features are typically extracted using matrices like the gray-level co-occurrence matrix (GLCM), gray-level size zone matrix (GLSZM), gray-level distance zone matrix (GLDZM), gray-level run-length matrix (GLRLM), and neighboring gray-tone difference matrix (NGTDM) (26).

Twenty-eight (28) radiomic feature-ROI pairs had significant associations with the SDMT scores in the Discovery study and met our threshold of significance in the replication analysis (FDR< 0.05) in which an independent set of 63 MS Snapshot participants were considered (Table 1). There were 19 unique shape, first-order or texture features within these 28 feature-ROI combinations. Thirteen (13) radiomic features derived from the whole thalamus, 9 from the pulvinar, 4 from the intralaminar and 2 from the medial nuclei.

**Table 1.**
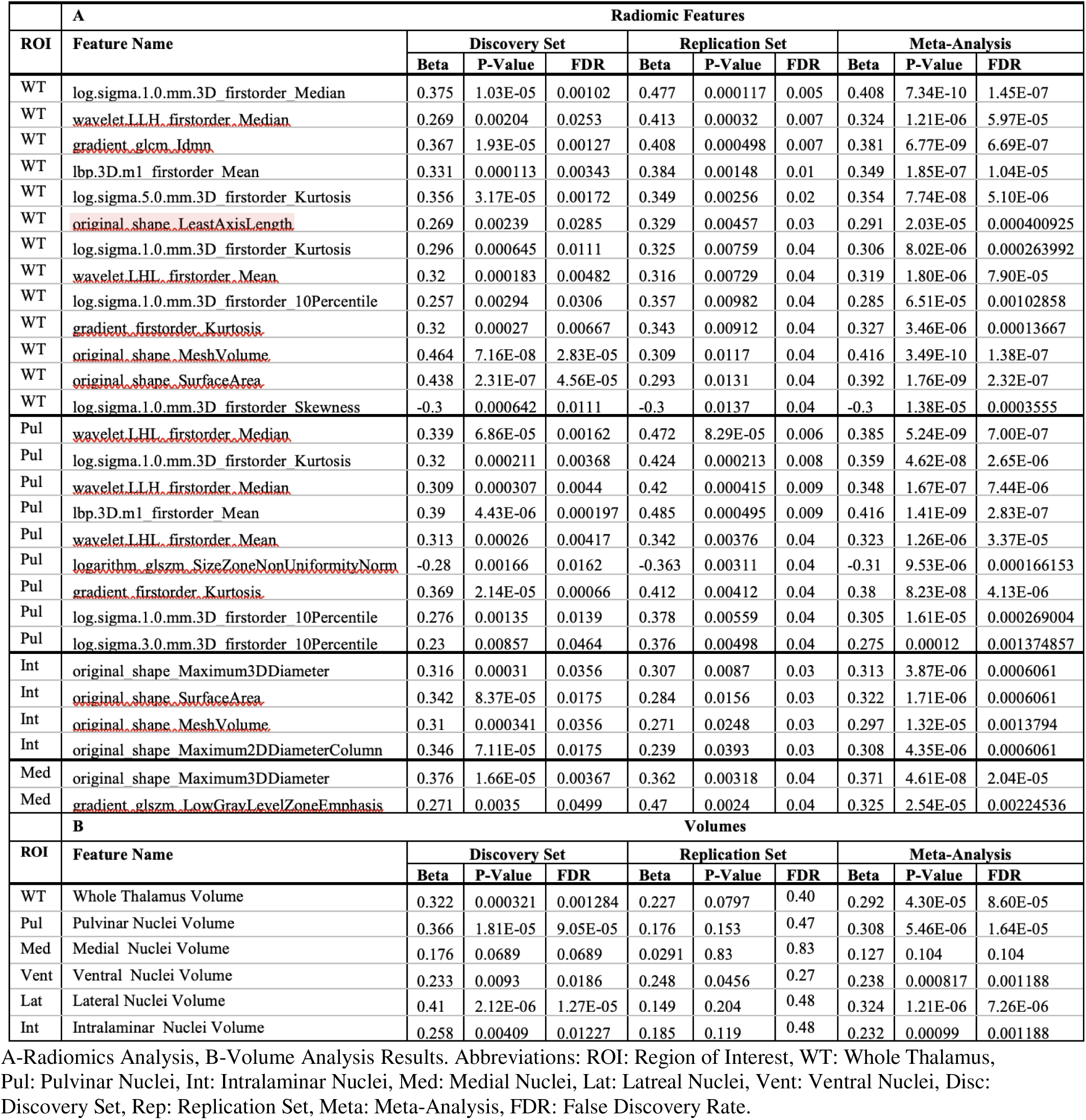
Discovery-Validation Set and Meta-Analysis Results.

The top 4 feature/ROI combinations were wavelet LHL.firstorder.Median (pulvinar nuclei, FDR= 0.006), log.sigma.1.0.mm.3D.first.order.Median (whole thalamus, FDR=0.005), original.shape.Maximum.3D.Diameter (medial nuclei, FDR=0.05) and original.shape.Maximum.3D.Diameter (intralaminar nuclei, FDR=0.03), illustrating the different types of features that reproducibly associated with SDMT in our data. A meta-analysis of the two datasets was then performed to provide the summary results from all available data (Table 1) to create a resource with summary evidence of the data.

In parallel, we repeated these analyses using traditional thalamic volume measures derived from the same ROIs to offer a comparison with what volumetric analyses would return from the same participants with MRI data. We used the same analytic method, adjusting for age and sex. The volumetric measures were significantly associated with SDMT in the Discovery study, but these associations did not replicate: we note that the volume of the ventral nuclei had a p-value of 0.045 in the replication analysis, yielding a corresponding FDR value of 0.27 (Table 1), indicating it did not survive correction for multiple comparisons. Thus, while our volumetric analyses capture the reported associations of thalamic volume with SDMT in PwMS, the associations with radiomic features appear to be more statistically robust.

To illustrate these results more clearly, Figure 3 A-B presents forest plots displaying the effect size and 95% confidence interval for the whole thalamus volume alongside its significant radiomic feature log.sigma.1.0.mm.3D.firstorder, across the discovery, replication, and meta-analysis phases, as well as for the pulvinar nuclei volume and its significant radiomic feature wavelet LHL.first.order.Median (Figure 3 C-D). The replicated radiomic features have a stronger effect size in their association with SDMT than the traditional volumetric measures.

**Figure 3.**
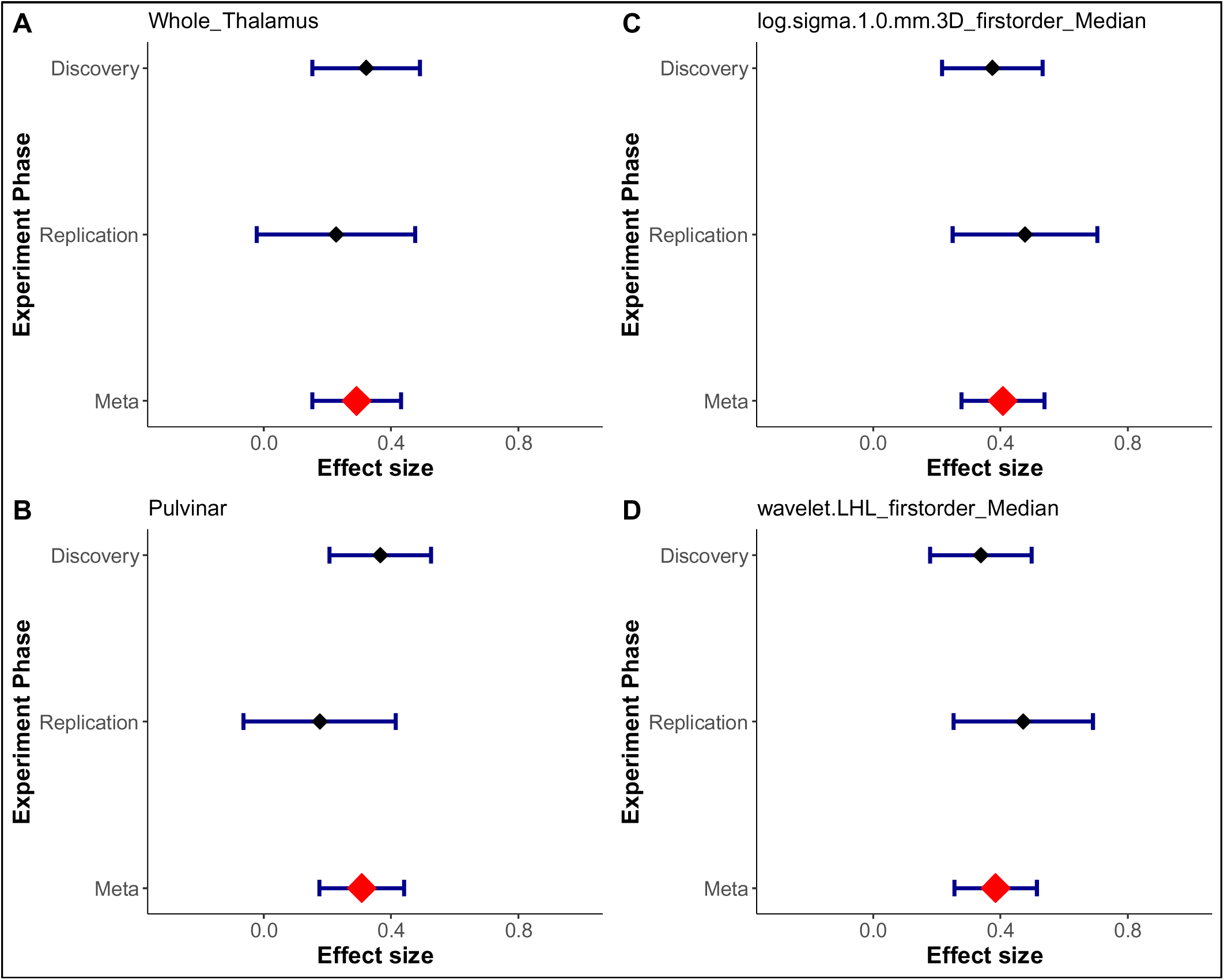
Forest plots showing effect sizes and 95% confidence intervals for whole thalamus volume and its most significant radiomic feature (log.sigma.1.0.mm.3D.firstorder), and for pulvinar volume and its most significant feature (wavelet.LHL.firstorder.Median) across discovery, replication, and meta-analysis phases. The replicated radiomic features showed stronger effect sizes in their association with the Symbol Digit Modalities Test than the volumetric measures.

### Putamen-Derived Radiomic Features-SDMT Associations

To demonstrate the specificity of our results to the thalamus, we performed a linear regression analysis controlling for age and sex, using the above mentioned 19 unique features derived from a different deep gray matter structure, the putamen. Only 4 out of the 19 unique features were found to be significantly associated with SDMT scores: original shape Mesh Volume (FDR=0.01), original shape Least Axis Length, log sigma 3.0 mm 3D first order 10^th^ Percentile, and original shape Surface Area (all FDR=0.04). These findings support the specificity of our results, suggesting that the observed radiomic changes in MS may be more prominent in the thalamus than in other structures, suggesting some specificity of our associations. The full list of putamen-derived radiomic features and their associations with SDMT is reported in Supplementary Table 2.

### Associations Between Significant Radiomic Features, PASAT and DSST Scores

In the second set of clinical MRI data collected by our colleagues at the Mayo Clinic (N=176) from PwMS who were assessed using the PASAT, we used the same linear regression model adjusted for age and sex to assess wheter we can replicate our results from Table 1 in a distinct set of individuals, with MRIs collected in a different protocol, and using a distinct cognitive test measuring the performance in similar cognitive domains. Of the 28 radiomic feature-ROI pairs replicated in the MS Snapshot analysis, 24 (12 from the whole thalamus, 7 from the pulvinar, 4 from the intralaminar, and 1 from the medial nuclei) were found to be associated with PASAT scores (FDR < 0.05) in the Mayo Clinic participants (Table 2).

**Table 2.**
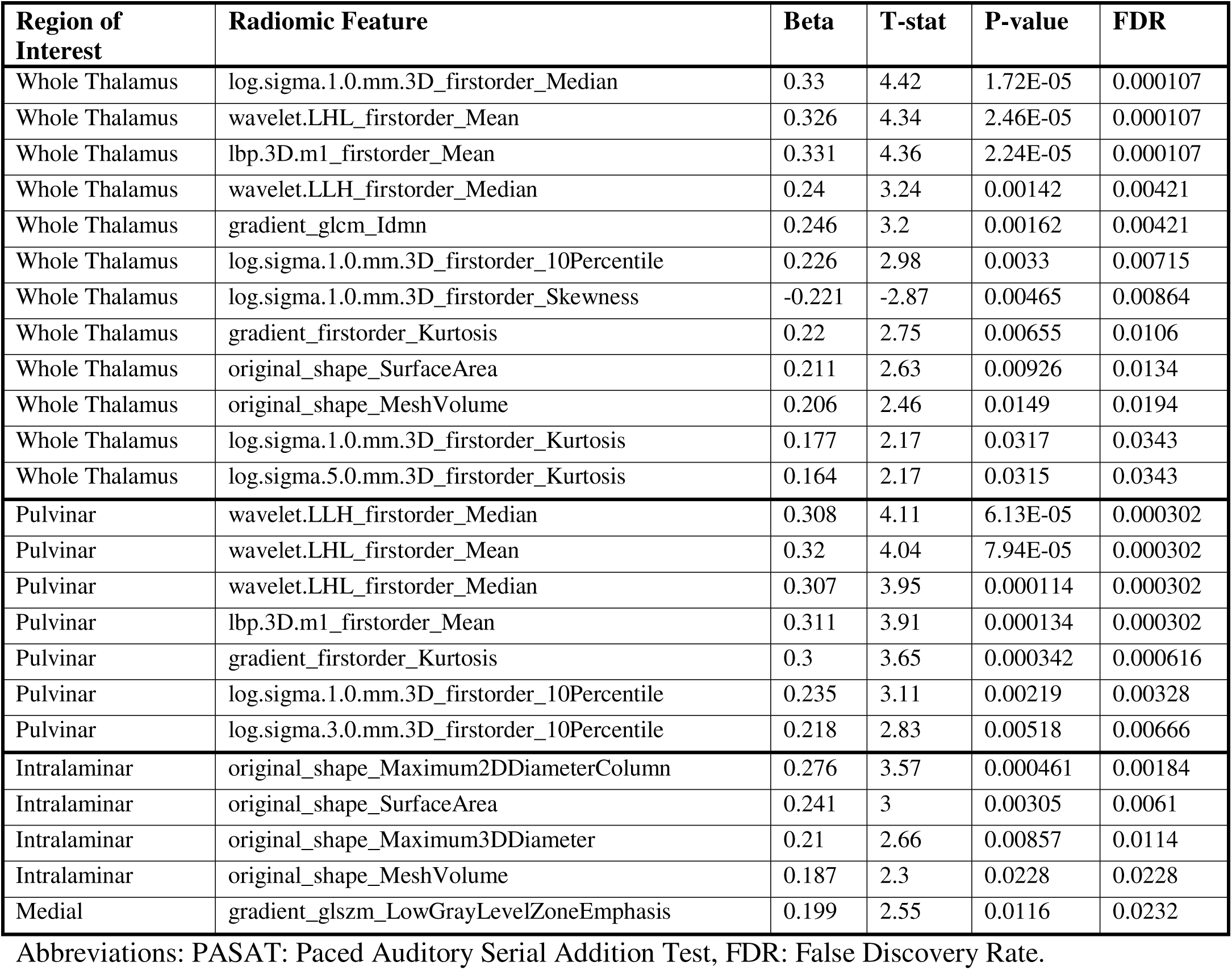
Radiomic features significantly predict PASAT Scores.

We then turned to the RANN participants (N=259), who do not have MS. Of the 28 radiomic feature-ROI pairs presented in Table 1, only 2 were associated with the DSST results in these individuals recruited from the general population to sample the lifespan: original.shape.Maximum.2D.Diameter.Column from the intralaminar nuclei (FDR=0.003) and original.shape.Surface.Area from the whole thalamus (FDR=0.027). Detailed results are available in Supplementary Table 3. These results suggest that our results from the MS Snapshot study are relatively specific to the context of MS, although aging-related processes may have an effect on two of the 28 radiomic features.

### LASSO Analysis Results

Given the discovery of replicable radiomic features associated with SDMT, we wanted to identify the combination of features that best predicted our measure of cognitive performance. We therefore deployed a LASSO method to predict SDMT. We developed three models: one from the radiomic features, one from the volumetric measures and one model combining the radiomic and volumetric features. The radiomic model had higher r^2^ values and lower p-values compared to volume-based models across each thalamic nucleus. For example, in pulvinar, radiomics achieved a median p-value of 0.0003 and an r^2^ of 0.23, while the volume-based model had a p-value of 0.005 and an r^2^ of 0.14. Among the predictions based on individual nuclei, the radiomics r^2^ was typically at least twice as large as the volumetric-based prediction. The best r^2^ was found in the medial nuclei where the radiomics models (p-value=0.002 and r^2^=0.17) outperformed the volumetric models (p-value=0.05 and r^2^=0.07). The results are shown in Table 3. Combining radiomic and volumetric features yielded better results than the volumetric-only models; however, it did not perform better than the radiomics-only model.

**Table 3.** Symbol Digit Modalities Test (SDMT) Prediction Results.

| Models | Whole Thalamus |  | Pulvinar |  | Medial |  | Ventral |  | Lateral |  | Intralaminar |  |
| --- | --- | --- | --- | --- | --- | --- | --- | --- | --- | --- | --- | --- |
|  | R<sup>2</sup> | P | R<sup>2</sup> | P | R<sup>2</sup> | P | R<sup>2</sup> | P | R | P | R<sup>2</sup> | P |
| Radiomics | 0.18 | 0.001 | 0.23 | 0.0003 | 0.17 | 0.002 | 0.17 | 0.002 | 0.18 | 0.001 | 0.11 | 0.01 |
| Volumes | 0.15 | 0.004 | 0.14 | 0.006 | 0.07 | 0.05 | 0.11 | 0.01 | 0.16 | 0.003 | 0.09 | 0.03 |
| Radiomics + Volumes | 0.16 | 0.003 | 0.23 | 0.0003 | 0.17 | 0.002 | 0.17 | 0.002 | 0.18 | 0.001 | 0.11 | 0.01 |

P and R<sup>2</sup> values represent median values of the 1000 simulations.

In the data derived from the healthy RANN study participants (N=259), we found the same pattern: models built solely on radiomic features predicted DSST results more accurately than models derived from volumetric measures only. None of the volumetric models showed significance, while all radiomic models except lateral nuclei were found significantly associated with DSST scores (p-values ranging from 0.009 to 0.05 and r^2^ from 0.05 to 0.08). Incorporating volumetric measures alongside radiomic features did not enhance prediction beyond the radiomics-only model (Table 4).

**Table 4.**
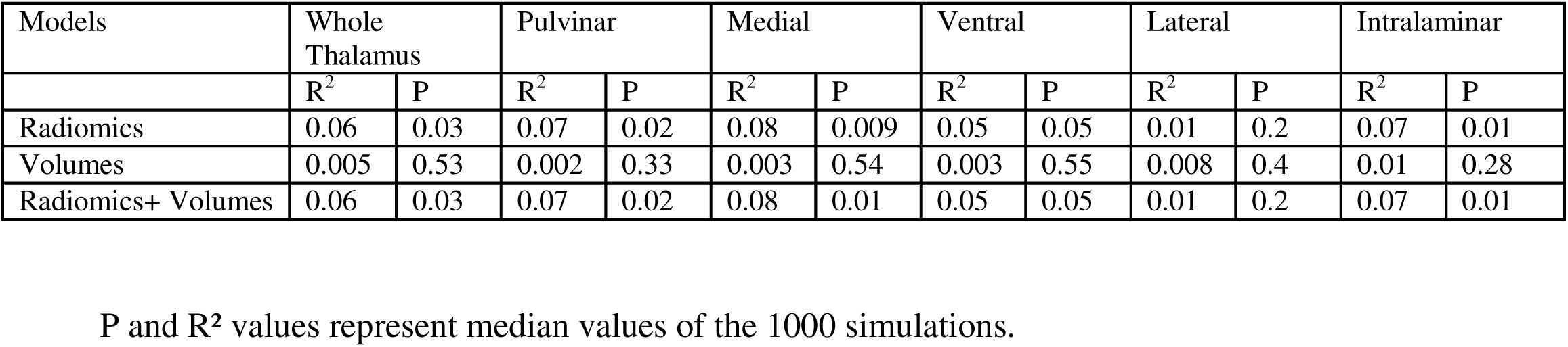
Digit Symbol Substitution Test Prediction Results.

## Discussion

Radiomics can capture spatial complexity and intraregional heterogeneity often overlooked by conventional volumetric analyses or visual assessment of structural MRI. Given its critical role in MS, we focused our analysis on the thalamus and aimed to identify radiomic features associated with cognitive performance. Importantly, our analyses were based on repurposed MRI data initially acquired during routine clinical care, highlighting the potential for implementing this radiomics-based approach in real-world clinical settings to inform patient management and prognosis.

To anchor our analyses, we selected the SDMT as the primary cognitive outcome, given its widespread use in PwMS and its well-established associations with thalamic abnormalities (38). Additionally, we included other measures of processing speed, such as the PASAT and the DSST, in independent cohorts to assess the generalizability and specificity of our findings.

Our primary analytic framework (discovery, replication, and meta-analysis) demonstrated that radiomic features were more strongly associated with SDMT performance in PwMS than conventional volumetric measures. While 28 radiomic feature-ROI pairs showed significant, replicated relationships, only the volume of the ventral thalamic nuclei was suggestively linked to SDMT scores in the replication set (Table 1).

Most of the radiomic features associated with the SDMT scores in this analysis were extracted from the whole thalamus (13) and the pulvinar (9). Twenty of these features (17 first order, 2 GLSZM and 1 GLCM features) capture the intensity distribution and spatial relationships between voxels, reflecting patterns, heterogeneity, or regularity the within ROIs. Generally, greater intensity variation within an ROI could suggest tissue damage, demyelination, or neurodegeneration, whereas more uniform textures may indicate areas that are preserved or less affected (14). The remaining 8 features were shape features, which quantify the geometric characteristics of a region including size, volume, surface area, sphericity, and complexity (47). These features were primarily concentrated in the intralaminar nuclei, likely reflecting the influence of the size of the region, as larger regions are more capable of capturing textural heterogeneity.

To validate the anatomical specificity of our findings, we conducted a control analysis using radiomic features derived from the putamen given its proximity and comparable size to the whole thalamus. Notably, only 4 of the putamen-derived features showed association with SDMT scores. This highlights the region-specific nature of our primary findings as well as underscores the functional relevance of thalamic radiomic abnormalities in cognitive performance.

When evaluated in an independent cohort of PwMS, 24 of the 28 radiomic feature-ROI pairs linked to SDMT performance also showed strong associations with PASAT outcomes underscoring the generalizability of our findings. The SDMT and PASAT likely engage different brain regions and networks as they asses the processing speed through distinct tasks. Nonetheless, our findings revealed a common radiomic signature in the thalamus that is highly significant across both measures. This underscores the robustness and generalizability of these features, reflecting consistent and biologically meaningful thalamic radiomic characteristics associated mainly with processing speed across different tests. Furthermore, these radiomic features did not predict DSST performance (a variant of the SDMT) in healthy controls. This finding supports the specificity of these features to MS.

Our observation was also robust to the analytic approach that was deployed, as illustrated by the results of our LASSO models, in which radiomic features outperformed volume-based models in predicting SDMT results across all thalamic nuclei as well (Table 3). LASSO analyses also showed that combining radiomic and volumetric features produced better models than those using volumetric features alone. However, they did not outperform models based solely on radiomics in contrary to similar studies (28).

In healthy participants, we found similar relative differences between the associations of the DSST with radiomic versus volumetric measures (Table 4). That is, although the important features were different for PwMS and RANN participants (Supplementary Tables 4-5), radiomics performed better than the volumetrics in both cohorts. This suggests that our approach is robust to the participant population and to different MRI protocols. The associations in healthy controls were attenuated, which is not surprising since these are cognitively non-impaired healthy participants and therefore have a narrower range of outcomes than PwMS.

In all analyses, the pulvinar emerged as particularly important nuclei in our study amongst others. Of the 28 repeated radiomic feature-ROI pairs in the discovery-replication analysis, 9 originated from the pulvinar, the highest count after the whole thalamus (13 features). Notably, 7 of these 9 pulvinar features were also associated with PASAT performance, highlighting their consistent cognitive relevance. In LASSO regression, the most significant radiomic predictors were also pulvinar-derived (R²=0.23, p=0.0003).

The pulvinar plays a critical role in visual attention and cognitive integration through its extensive connections with both cortical and subcortical regions (48–50). Structural and functional abnormalities of the pulvinar have been documented across various neurological disorders, including MS (51,52). Notably, a recent study reported iron depletion in the pulvinar in association with paramagnetic rim lesions, suggesting a potential link between microglial activity and iron loss in MS (53). Given its functional relevance and emerging pathological findings, a more targeted investigation of the pulvinar using radiomics may offer valuable insights into the mechanisms underlying cognitive deficits in MS.

Although the literature on thalamus and T1 derived radiomics is limited, our findings align with previous studies demonstrating the relevance of first-order and texture features in MS (54), Parkinson’s disease, and epilepsy (55,56). Similarly, associations between thalamic wavelet features and cognitive performance have been observed in subcortical ischemic vascular impairment (57). These consistent findings across neurological conditions suggest that radiomic features capturing voxel intensity and spatial patterns may serve as valuable markers of thalamic integrity, with potential relevance for understanding and predicting disease outcomes.

A key limitation of our study is the lack of uniform cognitive assessments across cohorts. While both the DSST and SDMT are highly sensitive in detecting processing speed, attention, and working memory, subtle differences in design could affect performance. Similarly, although both the SDMT and PASAT primarily assess cognitive processing speed, working memory, attention, concentration, and information processing they differ in modality: the SDMT is centered on visual scanning, while the PASAT focuses on auditory and arithmetic processing. Therefore, caution is needed when directly comparing cognitive performance measured by SDMT, DSST and PASAT.

Additionally, while our analysis focused primarily on processing speed, considering the other cognitive domains affected in MS future research should incorporate a broader range of cognitive assessments and regions of interest (such as the hippocampus) to better characterize the relationship between radiomic features and specific cognitive functions, including episodic memory.

We used only 3D T1-weighted images in this study which can be considered as another limitation. This was to ensure consistency across cohorts, as 3D FLAIR sequences were not available for all participants across different cohorts. Considering their essential role in detecting MS-related damage (i.e., lesions), incorporating sequences like T2 FLAIR could enhance the sensitivity and clinical relevance of radiomic analyses.

Although we observed strong associations between thalamus derived radiomic features which primarily reflect tissue intensity distribution, and cognitive performance, their biological relevance in MS remains uncertain. For radiomics to achieve clinical utility, it is essential to establish links between quantitative features and underlying biological processes, including molecular, metabolic, genetic, microstructural, and histopathological changes. Postmortem studies have shown that neurodegeneration and diffuse inflammation can affect normal-appearing thalamic gray matter in MS (4). These changes may not be detectable on conventional structural MRI. Therefore, investigating the extent to which radiomics can non-invasively capture these subtle pathological alterations represents a promising direction for future research. Given reports of iron abnormalities in the pulvinar, investigating whether radiomics can capture such changes on standard clinical images presents another promising avenue for further research. Ultimately, demonstrating the ability of radiomics to identify these microstructural alterations and integrating this information with other biological data, such as proteomics and genomics represents a critical step toward clinical translation.

In conclusion, our study demonstrated the potential of radiomics to extract clinically relevant information from conventional MRI scans, revealing aspects that traditional volumetric measurements in clinical MRI cannot capture. With further validation, radiomics could play a significant role in clinical practice by providing early biomarkers for cognitive deficits, enhancing the understanding of thalamic involvement in MS.

## Acknowledgements and Disclosure

We thank the participants of the MS Snapshot, RANN, and Mayo Clinic studies for their time and participation. We also thank Dr. Victoria M. Leavitt for developing the comprehensive cognitive test battery used in the MS Snapshot study and Columbia University MS Center Coordinators Orian Amona and Julia Goralsky for their assistance with data acquisition.

## Notes

### Competing Interest Statement

The authors have declared no competing interest.

## References

1. Benedict RHB, Amato MP, DeLuca J, Geurts JJG. Cognitive impairment in multiple sclerosis: clinical management, MRI, and therapeutic avenues. The Lancet Neurology. 2020 Oct 1;19(10):860–71.

2. Torrico TJ, Munakomi S. Neuroanatomy, Thalamus. In: StatPearls [Internet]. Treasure Island (FL): StatPearls Publishing; 2025 [cited 2025 Jan 8]. Available from: http://www.ncbi.nlm.nih.gov/books/NBK542184/

3. Roy DS, Zhang Y, Halassa MM, Feng G. Thalamic subnetworks as units of function. Nat Neurosci. 2022 Feb;25(2):140–53.

4. Kipp M, Wagenknecht N, Beyer C, Samer S, Wuerfel J, Nikoubashman O. Thalamus pathology in multiple sclerosis: from biology to clinical application. Cell Mol Life Sci. 2015 Mar;72(6):1127–47.

5. Benedict RH, Hulst HE, Bergsland N, Schoonheim MM, Dwyer MG, Weinstock-Guttman B, et al. Clinical significance of atrophy and white matter mean diffusivity within the thalamus of multiple sclerosis patients. Mult Scler. 2013 Oct;19(11):1478–84.

6. Muhlert N, Atzori M, De Vita E, Thomas DL, Samson RS, Wheeler-Kingshott CAM, et al. Memory in multiple sclerosis is linked to glutamate concentration in grey matter regions. Journal of Neurology, Neurosurgery & Psychiatry. 2014 Aug 1;85(8):833–9.

7. Derache N, Marié RM, Constans JM, Defer GL. Reduced thalamic and cerebellar rest metabolism in relapsing-remitting multiple sclerosis, a positron emission tomography study: correlations to lesion load. J Neurol Sci. 2006 Jun 15;245(1–2):103–9.

8. d’Ambrosio A, Hidalgo De La Cruz M, Valsasina P, Pagani E, Colombo B, Rodegher M, et al. Structural connectivity defined thalamic subregions have different functional connectivity abnormalities in multiple sclerosis patients: Implications for clinical correlations. Human Brain Mapping. 2017 Dec;38(12):6005–18.

9. Mahajan KR, Nakamura K, Cohen JA, Trapp BD, Ontaneda D. Intrinsic and Extrinsic Mechanisms of Thalamic Pathology in Multiple Sclerosis. Ann Neurol. 2020 Jul;88(1):81– 92.

10. Azevedo CJ, Cen SY, Khadka S, Liu S, Kornak J, Shi Y, et al. Thalamic atrophy in multiple sclerosis: A magnetic resonance imaging marker of neurodegeneration throughout disease. Ann Neurol. 2018 Feb;83(2):223–34.

11. Van Timmeren JE, Cester D, Tanadini-Lang S, Alkadhi H, Baessler B. Radiomics in medical imaging—“how-to” guide and critical reflection. Insights Imaging. 2020 Dec;11(1):91.

12. Lin FY, Chang YC, Huang HY, Li CC, Chen YC, Chen CM. A radiomics approach for lung nodule detection in thoracic CT images based on the dynamic patterns of morphological variation. Eur Radiol. 2022 Jun 1;32(6):3767–77.

13. Park YW, Choi YS, Kim SE, Choi D, Han K, Kim H, et al. Radiomics features of hippocampal regions in magnetic resonance imaging can differentiate medial temporal lobe epilepsy patients from healthy controls. Sci Rep. 2020 Nov 11;10(1):19567.

14. Zhang Y, Zhu H, Mitchell JR, Costello F, Metz LM. T2 MRI texture analysis is a sensitive measure of tissue injury and recovery resulting from acute inflammatory lesions in multiple sclerosis. NeuroImage. 2009 Aug 1;47(1):107–11.

15. Lu L, Dercle L, Zhao B, Schwartz LH. Deep learning for the prediction of early on-treatment response in metastatic colorectal cancer from serial medical imaging. Nat Commun. 2021 Nov 17;12(1):6654.

16. Dercle L, Fronheiser M, Lu L, Du S, Hayes W, Leung DK, et al. Identification of Non–Small Cell Lung Cancer Sensitive to Systemic Cancer Therapies Using Radiomics. Clinical Cancer Research. 2020 May 1;26(9):2151–62.

17. Lu L, Sun SH, Yang H, E L, Guo P, Schwartz LH, et al. Radiomics Prediction of EGFR Status in Lung Cancer-Our Experience in Using Multiple Feature Extractors and The Cancer Imaging Archive Data. Tomography. 2020 Jun 1;6(2):223–30.

18. Van Griethuysen JJM, Fedorov A, Parmar C, Hosny A, Aucoin N, Narayan V, et al. Computational Radiomics System to Decode the Radiographic Phenotype. Cancer Research. 2017 Nov 1;77(21):e104–7.

19. Tupe-Waghmare P, Rajan A, Prasad S, Saini J, Pal PK, Ingalhalikar M. Radiomics on routine T1-weighted MRI can delineate Parkinson’s disease from multiple system atrophy and progressive supranuclear palsy. Eur Radiol. 2021 Nov;31(11):8218–27.

20. Feng Q, Ding Z. MRI Radiomics Classification and Prediction in Alzheimer’s Disease and Mild Cognitive Impairment: A Review. CAR. 2020 May 18;17(3):297–309.

21. Ma X, Zhang L, Huang D, Lyu J, Fang M, Hu J, et al. Quantitative radiomic biomarkers for discrimination between neuromyelitis optica spectrum disorder and multiple sclerosis. Magnetic Resonance Imaging. 2019 Apr;49(4):1113–21.

22. Huang J, Xin B, Wang X, Qi Z, Dong H, Li K, et al. Multi-parametric MRI phenotype with trustworthy machine learning for differentiating CNS demyelinating diseases. J Transl Med. 2021 Dec;19(1):377.

23. He T, Zhao W, Mao Y, Wang Y, Wang L, Kuang Q, et al. MS or not MS: T2-weighted imaging (T2WI)-based radiomic findings distinguish MS from its mimics. Mult Scler Relat Disord. 2022 May;61:103756.

24. Peng Y, Zheng Y, Tan Z, Liu J, Xiang Y, Liu H, et al. Prediction of unenhanced lesion evolution in multiple sclerosis using radiomics-based models: a machine learning approach. Multiple Sclerosis and Related Disorders. 2021 Aug;53:102989.

25. Wang X, Wang X, Xu Y, Yan Z, Shi Z, Liu Y, et al. Radiomics-based analysis of choroid plexus abnormalities in neuromyelitis optica spectrum disorders and multiple sclerosis and their clinical implications. Mult Scler Relat Disord. 2025 Apr 22;99:106465.

26. Inglese M, Conti A, Toschi N. Radiomics across modalities: a comprehensive review of neurodegenerative diseases. Clinical Radiology. 2025 Jun 1;85:106921.

27. Yan Z, Yuan S, Zhu Q, Wang X, Shi Z, Zhang Y, et al. Radiomics models based on cortical damages for identification of multiple sclerosis with cognitive impairment. Multiple Sclerosis and Related Disorders. 2024 Jan;81:105348.

28. Wang X, Liu S, Yan Z, Yin F, Feng J, Liu H, et al. Radiomics Nomograms Based on Multi-sequence MRI for Identifying Cognitive Impairment and Predicting Cognitive Progression in Relapsing-Remitting Multiple Sclerosis. Acad Radiol. 2024 Aug 27;S1076-6332(24)00591–9.

29. Chiaravalloti ND, DeLuca J. Cognitive impairment in multiple sclerosis. Lancet Neurol. 2008 Dec;7(12):1139–51.

30. Strober L, DeLuca J, Benedict RH, Jacobs A, Cohen JA, Chiaravalloti N, et al. Symbol Digit Modalities Test: A valid clinical trial endpoint for measuring cognition in multiple sclerosis. Mult Scler. 2019 Nov;25(13):1781–90.

31. Buyukturkoglu K, Dworkin JD, Leiva V, Provenzano FA, Guevara P, De Jager PL, et al. Brain volumetric correlates of remotely versus in-person administered symbol digit modalities test in multiple sclerosis. Multiple Sclerosis and Related Disorders. 2022 Dec;68:104247.

32. Ghandili M, Munakomi S. Neuroanatomy, Putamen. In: StatPearls [Internet]. Treasure Island (FL): StatPearls Publishing; 2025 [cited 2025 May 16]. Available from: http://www.ncbi.nlm.nih.gov/books/NBK542170/

33. Simani L, Molaeipour L, Kian S, Leavitt VM. Correlation between cognitive changes and neuroradiological changes over time in multiple sclerosis: a systematic review and meta-analysis. J Neurol. 2024 Aug;271(8):5498–518.

34. Zeydan B, Son J, Neyal N, Schwarz CG, Atkinson EJ, Morrison HA, et al. Upper cervical spinal cord atrophy in MS: Sex, menopause, and neurodegeneration. Mult Scler. 2025 Jan 18;13524585241311441.

35. Stern Y, Habeck C, Steffener J, Barulli D, Gazes Y, Razlighi Q, et al. The Reference Ability Neural Network Study: Motivation, design, and initial feasibility analyses. NeuroImage. 2014 Dec;103:139–51.

36. Jaeger J. Digit Symbol Substitution Test: The Case for Sensitivity Over Specificity in Neuropsychological Testing. J Clin Psychopharmacol. 2018 Oct;38(5):513–9.

37. Smith A. Symbol Digit Modalities Test (SDMT). Manual (Revised). Los Angeles: Western Psychological Services.; 1982.

38. Rao SM, Martin AL, Huelin R, Wissinger E, Khankhel Z, Kim E, et al. Correlations between MRI and Information Processing Speed in MS: A Meta-Analysis. Multiple Sclerosis International. 2014;2014:1–9.

39. Levy S, Dvorak EM, Graney R, Staker E, Sumowski JF. In-person and remote administrations of the symbol digit modalities test are interchangeable among persons with multiple sclerosis. Multiple Sclerosis and Related Disorders. 2023 Mar 1;71:104553.

40. Rao SM, Leo GJ, Ellington L, Nauertz T, Bernardin L, Unverzagt F. Cognitive dysfunction in multiple sclerosis.: II. Impact on employment and social functioning. Neurology. 1991 May;41(5):692–6.

41. Buyukturkoglu K, Mormina E, De Jager PL, Riley CS, Leavitt VM. The Impact of MRI T1 Hypointense Brain Lesions on Cerebral Deep Gray Matter Volume Measures in Multiple Sclerosis. J Neuroimaging. 2019 Jul;29(4):458–62.

42. González-Villà S, Valverde S, Cabezas M, Pareto D, Vilanova JC, Ramió-Torrentà L, et al. Evaluating the effect of multiple sclerosis lesions on automatic brain structure segmentation. Neuroimage Clin. 2017;15:228–38.

43. Cerri S, Puonti O, Meier DS, Wuerfel J, Mühlau M, Siebner HR, et al. A contrast-adaptive method for simultaneous whole-brain and lesion segmentation in multiple sclerosis. NeuroImage. 2021 Jan 15;225:117471.

44. Iglesias JE, Insausti R, Lerma-Usabiaga G, Bocchetta M, Van Leemput K, Greve DN, et al. A probabilistic atlas of the human thalamic nuclei combining ex vivo MRI and histology. NeuroImage. 2018 Dec;183:314–26.

45. Orlhac F, Eertink JJ, Cottereau AS, Zijlstra JM, Thieblemont C, Meignan M, et al. A Guide to ComBat Harmonization of Imaging Biomarkers in Multicenter Studies. J Nucl Med. 2022 Feb;63(2):172–9.

46. Tibshirani R. Regression Shrinkage and Selection via the Lasso. Journal of the Royal Statistical Society Series B (Methodological). 1996;58(1):267–88.

47. Tomaszewski MR, Gillies RJ. The Biological Meaning of Radiomic Features. Radiology. 2021 Mar;298(3):505–16.

48. Grieve KL, Acuña C, Cudeiro J. The primate pulvinar nuclei: vision and action. Trends in Neurosciences. 2000 Jan;23(1):35–9.

49. LaBerge D, Buchsbaum MS. Positron emission tomographic measurements of pulvinar activity during an attention task. J Neurosci. 1990 Feb;10(2):613–9.

50. Fama R, Sullivan EV. Thalamic structures and associated cognitive functions: Relations with age and aging. Neuroscience & Biobehavioral Reviews. 2015 Jul;54:29–37.

51. Fadda L, Floris G, Polizzi L, Meleddu L, Ercoli T, Garofalo P, et al. Pulvinar sign in a case of anti-CV2 encephalitis. J Neurol Sci. 2018 Oct 15;393:69–71.

52. Trufanov A, Bisaga G, Skulyabin D, Temniy A, Poplyak M, Chakchir O, et al. Thalamic nuclei degeneration in multiple sclerosis. Journal of Clinical Neuroscience. 2021 Jul;89:375– 80.

53. Reeves JA, Salman F, Mohebbi M, Bergsland N, Jakimovski D, Hametner S, et al. Association between paramagnetic rim lesions and pulvinar iron depletion in persons with multiple sclerosis. Multiple Sclerosis and Related Disorders. 2025 Jan 1;93:106187.

54. Pontillo G, Tommasin S, Cuocolo R, Petracca M, Petsas N, Ugga L, et al. A Combined Radiomics and Machine Learning Approach to Overcome the Clinicoradiologic Paradox in Multiple Sclerosis. AJNR Am J Neuroradiol. 2021 Nov;42(11):1927–33.

55. Tupe-Waghmare P, Rajan A, Prasad S, Saini J, Pal PK, Ingalhalikar M. Radiomics on routine T1-weighted MRI can delineate Parkinson’s disease from multiple system atrophy and progressive supranuclear palsy. Eur Radiol. 2021 Nov 1;31(11):8218–27.

56. Sim Y, Lee SK, Chu MK, Kim WJ, Heo K, Kim KM, et al. MRI-Based Radiomics Approach for Differentiating Juvenile Myoclonic Epilepsy from Epilepsy with Generalized Tonic-Clonic Seizures Alone. J Magn Reson Imaging. 2024 Jul;60(1):281–8.

57. Liu B, Meng S, Cheng J, Zeng Y, Zhou D, Deng X, et al. Diagnosis of Subcortical Ischemic Vascular Cognitive Impairment With No Dementia Using Radiomics of Cerebral Cortex and Subcortical Nuclei in High-Resolution T1-Weighted MR Imaging. Front Oncol [Internet]. 2022 Apr 8 [cited 2025 May 13];12.

