## Supplementary Material for "A Radiomic Approach to Clinical MRI Refines the Thalamus-Cognition Link in Multiple Sclerosis"

**1- Demographic and cognitive variables**.

Supplementary Table 1.

| Participants | Age | Sex | Edu. | WTAR (Raw) | SDMT | DSST | PASAT |
| --- | --- | --- | --- | --- | --- | --- | --- |
| MS SNAPSHOT (N=168) | 48.5 ± 13 | 115 F | 16 ± 2 | 40 ± 9 | 43 ± 14 | N.A. | N.A. |
| RANN (N=259) | 52 ± 16 | 146 F | 16 ± 2 | 39 ± 9 | N.A. | 77 ± 14 | N.A. |
| MAYO Clinic (N=176) | 48 ± 13 | 127 F | N.A. | N.A. | N.A. | N.A. | 44 ± 12 |

All values reported as mean ± standard deviation. Abbreviations: WTAR: Wechsler Test of Adult Reading, SDMT: Symbol Digit Modalities Test, PASAT: Paced Auditory Serial Addition Test, DSST: Digit Symbol Substitution Test, F: Female, Edu: Years of Education, N.A.: Not Applicable.

**2- MRI Details**

For all cohorts, 3D T1-weighted images were acquired with 1 mm isotropic voxel resolution.

**Snapshot Cohort:** The majority of images (N = 113) were acquired on a 3T GE scanner (GE Healthcare, Wauwatosa, WI) using the following parameters: TE = 3.05 ms, TR = 7.38 ms, and flip angle = 13°. Additional data were acquired on scanners from other centers. The remaining 53 participants were scanned on Siemens scanners, either 3T (N = 37) or 1.5T (N = 16).

**Mayo Cohort**: Most scans (N = 111) were obtained on a 3T Siemens scanner (Siemens Healthineers, Germany) with parameters: TE = 2.9 ms, TR = 2.3 ms and flip angle = 9°. The remaining 65 participants were scanned on 3T GE scanners.

**RANN Study:** Imaging was performed on a 3T Philips Achieva system. Participants underwent a T1-weighted MPRAGE scan with TE = 3 ms, TR = 6.5 ms and flip angle = 8°.

**2- LASSO Analysis**

**Feature Selection**

The Least Absolute Shrinkage and Selection Operator (LASSO) regression algorithm was utilized to select important radiomics features. LASSO regression integrates an L1 regularization term into the linear regression cost function, proportional to the absolute values of the model's coefficients.^1^ This L1 regularization encourages sparsity in the linear model by setting some coefficients to zero, enabling LASSO to automatically select relevant features by eliminating those with zero coefficients. This method is especially useful for datasets with many features, as it reduces dimensionality and mitigates the risk of model overfitting. In this implementation, the L1 regularization term is defined as λ * Σ|β_i|, where λ is the regularization parameter controlling model sparsity, and β_i are the linear regression model's coefficients. The top 10 radiomics features were used in the predictive models

**Building Predictive Models**

We built several models to predict cognitive test outcomes (i.e., SDMT and DSST results), including: 1- Models using only radiomic features, 2- Only individual thalamic nuclei volumes, 3- Models using radiomic and volumetric features together.

For each set of radiomics models and cognitive outcomes the following steps were taken to estimate their R² and p-values of association:

1. Randomly assign 70% of the sample to the training dataset and 30% to the testing dataset.
2. Remove all independent variables with a standard deviation of 0 and scale the rest.
3. Use the "glmnet" package in R version 4.2.2 to run a 5-fold cross-validated LASSO regression on the training data, producing a predictive model of the outcome variable based on the independent variables.
4. Apply the model built with the training dataset to the testing dataset to generate predicted values of the outcome variable.
5. Calculate the Pearson correlation coefficient between the predicted and observed values in the testing dataset and report the corresponding p-value and R².
6. To assess the typical model performance and stabilize the R² and p-value measures, iterate steps 1-5 a total of 1,000 times to determine the median R² and p-value of association.

**Supplementary Table 2. Putamen Derived Radiomic Features’ SDMT Prediction Performance.**

| Radiomic Feature | beta | se | tstat | pval | fdr |
| --- | --- | --- | --- | --- | --- |
| original_shape_MeshVolume | 0.281 | 0.0728 | 3.86 | 0.000161 | 0.0119 |
| original_shape_LeastAxisLength | 0.251 | 0.072 | 3.49 | 0.000618 | 0.037 |
| log.sigma.3.0.mm.3D_firstorder_10Percentile | 0.24 | 0.069 | 3.48 | 0.000643 | 0.0372 |
| original_shape_SurfaceArea | 0.249 | 0.0718 | 3.47 | 0.000667 | 0.0373 |
| original_shape_Maximum2DDiameterColumn | 0.187 | 0.0706 | 2.66 | 0.00869 | 0.239 |
| original_shape_Maximum3DDiameter | 0.16 | 0.073 | 2.19 | 0.03 | 0.262 |
| wavelet.LLH_firstorder_Median | 0.151 | 0.0692 | 2.18 | 0.0308 | 0.262 |
| log.sigma.1.0.mm.3D_firstorder_Median | 0.143 | 0.0693 | 2.06 | 0.0408 | 0.264 |
| log.sigma.1.0.mm.3D_firstorder_10Percentile | 0.0788 | 0.0709 | 1.11 | 0.268 | 0.576 |
| gradient_glcm_Idmn | 0.0762 | 0.07 | 1.09 | 0.278 | 0.586 |
| log.sigma.5.0.mm.3D_firstorder_Kurtosis | -0.0632 | 0.07 | -0.903 | 0.368 | 0.688 |
| logarithm_glszm_SizeZoneNonUniformityNormalized | -0.058 | 0.07 | -0.828 | 0.409 | 0.73 |
| lbp.3D.m1_firstorder_Mean | 0.0337 | 0.0703 | 0.48 | 0.632 | 0.846 |
| wavelet.LHL_firstorder_Median | 0.0335 | 0.0702 | 0.478 | 0.634 | 0.848 |
| log.sigma.1.0.mm.3D_firstorder_Skewness | -0.0308 | 0.0708 | -0.436 | 0.664 | 0.866 |
| gradient_firstorder_Kurtosis | 0.0265 | 0.0702 | 0.377 | 0.706 | 0.888 |
| gradient_glszm_LowGrayLevelZoneEmphasis | 0.0206 | 0.0704 | 0.293 | 0.77 | 0.914 |
| log.sigma.1.0.mm.3D_firstorder_Kurtosis | -0.0185 | 0.0702 | -0.263 | 0.793 | 0.919 |
| wavelet.LHL_firstorder_Mean | 0.0155 | 0.0702 | 0.221 | 0.825 | 0.936 |

**Supplementary Table 3.** Associations Between Replicated Radiomic Features and DSST Performance.

| Region of Interest | Radiomic Feature | beta | se | tstat | pval | fdr |
| --- | --- | --- | --- | --- | --- | --- |
| Intralaminar | original_shape_Maximum2DDiameterColumn | 0.214 | 0.0641 | 3.34 | 0.000954 | 0.00382 |
| Whole_Thalamus | original_shape_SurfaceArea | 0.22 | 0.0708 | 3.11 | 0.00207 | 0.0269 |
| Whole_Thalamus | original_shape_MeshVolume | 0.185 | 0.0698 | 2.66 | 0.00839 | 0.0545 |
| Whole_Thalamus | log.sigma.5.0.mm.3D_firstorder_Kurtosis | -0.141 | 0.0602 | -2.34 | 0.02 | 0.0867 |
| Intralaminar | original_shape_Maximum3DDiameter | 0.124 | 0.062 | 1.99 | 0.0474 | 0.0948 |
| Intralaminar | original_shape_SurfaceArea | 0.108 | 0.0632 | 1.71 | 0.0882 | 0.118 |
| Whole_Thalamus | lbp.3D.m1_firstorder_Mean | 0.124 | 0.0588 | 2.1 | 0.0365 | 0.119 |
| Whole_Thalamus | original_shape_LeastAxisLength | 0.123 | 0.064 | 1.93 | 0.0551 | 0.143 |
| Pulvinar | logarithm_glszm_SizeZoneNonUniformityNormalized | 0.143 | 0.0597 | 2.39 | 0.0174 | 0.157 |
| Pulvinar | gradient_firstorder_Kurtosis | 0.12 | 0.059 | 2.03 | 0.0431 | 0.194 |
| Whole_Thalamus | log.sigma.1.0.mm.3D_firstorder_Median | 0.093 | 0.0592 | 1.57 | 0.118 | 0.256 |
| Intralaminar | original_shape_MeshVolume | 0.0681 | 0.0615 | 1.11 | 0.269 | 0.269 |
| Pulvinar | log.sigma.1.0.mm.3D_firstorder_Kurtosis | 0.0869 | 0.0592 | 1.47 | 0.144 | 0.324 |
| Pulvinar | wavelet.LLH_firstorder_Median | 0.0861 | 0.0587 | 1.47 | 0.144 | 0.324 |
| Medial | gradient_glszm_LowGrayLevelZoneEmphasis | 0.0762 | 0.0587 | 1.3 | 0.196 | 0.392 |
| Pulvinar | lbp.3D.m1_firstorder_Mean | 0.069 | 0.0595 | 1.16 | 0.248 | 0.446 |
| Medial | original_shape_Maximum3DDiameter | 0.0357 | 0.0685 | 0.521 | 0.603 | 0.603 |
| Whole_Thalamus | log.sigma.1.0.mm.3D_firstorder_Kurtosis | 0.0387 | 0.062 | 0.625 | 0.533 | 0.848 |
| Whole_Thalamus | gradient_firstorder_Kurtosis | 0.0325 | 0.0598 | 0.544 | 0.587 | 0.848 |
| Whole_Thalamus | gradient_glcm_Idmn | 0.0425 | 0.0618 | 0.688 | 0.492 | 0.848 |
| Whole_Thalamus | wavelet.LLH_firstorder_Median | 0.0254 | 0.0589 | 0.431 | 0.667 | 0.867 |
| Whole_Thalamus | log.sigma.1.0.mm.3D_firstorder_10Percentile | 0.013 | 0.0633 | 0.206 | 0.837 | 0.912 |
| Whole_Thalamus | wavelet.LHL_firstorder_Mean | 0.012 | 0.0601 | 0.199 | 0.842 | 0.912 |
| Whole_Thalamus | log.sigma.1.0.mm.3D_firstorder_Skewness | -0.0046 | 0.0639 | -0.072 | 0.943 | 0.943 |
| Pulvinar | log.sigma.1.0.mm.3D_firstorder_10Percentile | -0.0197 | 0.0629 | -0.313 | 0.754 | 0.96 |
| Pulvinar | log.sigma.3.0.mm.3D_firstorder_10Percentile | -0.0269 | 0.0621 | -0.432 | 0.666 | 0.96 |
| Pulvinar | wavelet.LHL_firstorder_Mean | -0.0048 | 0.0602 | -0.08 | 0.936 | 0.96 |
| Pulvinar | wavelet.LHL_firstorder_Median | -0.0030 | 0.0599 | -0.050 | 0.96 | 0.96 |

**Supplementary Table 4.** 10 Most Important Radiomic Features Predicting the Symbol Digit Modalities Test (SDMT) results. P and R² values represent median values.

| **Region of Interest** | **10 Most Important Features as Selected from Lasso Regression** | **Median R^2^** | **Median p value** |
| --- | --- | --- | --- |
| Whole_Thalamus | log.sigma.3.0.mm.3D_glcm_MaximumProbability | 0.22 | 0.0004 |
|  | log.sigma.5.0.mm.3D_firstorder_TotalEnergy |  |  |
|  | log.sigma.1.0.mm.3D_firstorder_Skewness |  |  |
|  | original_shape_MajorAxisLength |  |  |
|  | wavelet.LLH_gldm_LargeDependenceHighGrayLevelEmphasis |  |  |
|  | log.sigma.5.0.mm.3D_firstorder_90Percentile |  |  |
|  | exponential_firstorder_TotalEnergy |  |  |
|  | gradient_firstorder_Kurtosis |  |  |
|  | log.sigma.5.0.mm.3D_firstorder_Kurtosis |  |  |
|  | log.sigma.3.0.mm.3D_firstorder_Kurtosis |  |  |
| Pulvinar | log.sigma.5.0.mm.3D_firstorder_TotalEnergy | 0.23 | 0.0003 |
|  | log.sigma.5.0.mm.3D_firstorder_90Percentile |  |  |
|  | log.sigma.1.0.mm.3D_firstorder_Kurtosis |  |  |
|  | log.sigma.5.0.mm.3D_firstorder_Skewness |  |  |
|  | original_shape_Maximum2DDiameterRow |  |  |
|  | log.sigma.1.0.mm.3D_firstorder_Skewness |  |  |
|  | wavelet.LHH_firstorder_Kurtosis |  |  |
|  | log.sigma.5.0.mm.3D_gldm_DependenceVariance |  |  |
|  | gradient_firstorder_Kurtosis |  |  |
|  | logarithm_glszm_LargeAreaLowGrayLevelEmphasis |  |  |
| Medial | original_shape_Maximum3DDiameter | 0.18 | 0.002 |
|  | log.sigma.3.0.mm.3D_firstorder_Kurtosis |  |  |
|  | log.sigma.3.0.mm.3D_glcm_MaximumProbability |  |  |
|  | exponential_firstorder_TotalEnergy |  |  |
|  | log.sigma.3.0.mm.3D_gldm_DependenceVariance |  |  |
|  | original_shape_Maximum2DDiameterRow |  |  |
|  | square_glszm_LowGrayLevelZoneEmphasis |  |  |
|  | logarithm_glszm_GrayLevelNonUniformityNormalized |  |  |
|  | wavelet.HLH_firstorder_RootMeanSquared |  |  |
|  | log.sigma.3.0.mm.3D_glszm_GrayLevelNonUniformity |  |  |
| Ventral | log.sigma.3.0.mm.3D_glcm_MaximumProbability | 0.21 | 0.0007 |
|  | original_shape_SurfaceArea |  |  |
|  | log.sigma.5.0.mm.3D_firstorder_Maximum |  |  |
|  | original_shape_Maximum2DDiameterRow |  |  |
|  | log.sigma.3.0.mm.3D_firstorder_Skewness |  |  |
|  | wavelet.LLH_gldm_LargeDependenceHighGrayLevelEmphasis |  |  |
|  | log.sigma.3.0.mm.3D_firstorder_Kurtosis |  |  |
|  | original_shape_Maximum2DDiameterSlice |  |  |
|  | wavelet.LHH_firstorder_Kurtosis |  |  |
|  | original_shape_Sphericity |  |  |
| Lateral | original_shape_Maximum2DDiameterSlice | 0.19 | 0.001 |
|  | original_shape_LeastAxisLength |  |  |
|  | log.sigma.5.0.mm.3D_firstorder_Maximum |  |  |
|  | original_glcm_ClusterShade |  |  |
|  | log.sigma.5.0.mm.3D_glszm_LargeAreaLowGrayLevelEmphasis |  |  |
|  | wavelet.LLL_glszm_LargeAreaLowGrayLevelEmphasis |  |  |
|  | log.sigma.1.0.mm.3D_firstorder_InterquartileRange |  |  |
|  | wavelet.LHL_glcm_InverseVariance |  |  |
|  | original_shape_Maximum2DDiameterRow |  |  |
|  | wavelet.LLH_firstorder_RootMeanSquared |  |  |
| Intralaminar | original_shape_Maximum2DDiameterColumn | 0.08 | 0.03 |
|  | original_shape_Maximum2DDiameterRow |  |  |
|  | log.sigma.3.0.mm.3D_glszm_GrayLevelNonUniformityNormalized |  |  |
|  | original_shape_Maximum3DDiameter |  |  |
|  | original_shape_SurfaceArea |  |  |
|  | log.sigma.5.0.mm.3D_firstorder_TotalEnergy |  |  |
|  | exponential_firstorder_TotalEnergy |  |  |
|  | wavelet.LHL_firstorder_RootMeanSquared |  |  |
|  | gradient_glcm_ClusterShade |  |  |
|  | log.sigma.5.0.mm.3D_firstorder_Median |  |  |

**Supplementary Table 5.** 10 Most Important Radiomic Features Predicting the Digit Symbol Substitution Test (DSST) results. P and R² values represent median values.

| **Region of Interest** | **10 Most Important Features as Selected from Lasso Regression** | **Median R^2^** | **Median p value** |
| --- | --- | --- | --- |
| Whole_Thalamus | log.sigma.3.0.mm.3D_gldm_SmallDependenceLowGrayLevelEmphasis (700) | 0.06 | 0.03 |
|  | original_shape_SurfaceArea (560) |  |  |
|  | original_shape_MajorAxisLength (441) |  |  |
|  | original_shape_Sphericity (394) |  |  |
|  | log.sigma.5.0.mm.3D_glszm_ZonePercentage (346) |  |  |
|  | original_firstorder_Maximum (325) |  |  |
|  | wavelet.HLL_gldm_SmallDependenceLowGrayLevelEmphasis (241) |  |  |
|  | original_shape_Maximum3DDiameter (240) |  |  |
|  | lbp.3D.m1_glrlm_RunLengthNonUniformity (184) |  |  |
|  | wavelet.HHH_glszm_LargeAreaHighGrayLevelEmphasis (177) |  |  |
| Pulvinar | original_shape_SurfaceVolumeRatio (974) | 0.07 | 0.02 |
|  | log.sigma.5.0.mm.3D_gldm_SmallDependenceLowGrayLevelEmphasis (492) |  |  |
|  | log.sigma.3.0.mm.3D_glcm_Idmn (439) |  |  |
|  | lbp.3D.m1_glszm_GrayLevelNonUniformity (350) |  |  |
|  | original_shape_MeshVolume (257) |  |  |
|  | log.sigma.3.0.mm.3D_firstorder_Kurtosis (242) |  |  |
|  | original_shape_LeastAxisLength (222) |  |  |
|  | logarithm_firstorder_Maximum (185) |  |  |
|  | log.sigma.3.0.mm.3D_glcm_Idn (163) |  |  |
|  | log.sigma.5.0.mm.3D_glrlm_LongRunLowGrayLevelEmphasis (160) |  |  |
| Medial | lbp.3D.k_glrlm_GrayLevelNonUniformity (738) | 0.08 | 0.01 |
|  | original_shape_MajorAxisLength (487) |  |  |
|  | lbp.3D.m1_glrlm_RunLengthNonUniformity (471) |  |  |
|  | original_shape_MinorAxisLength (238) |  |  |
|  | original_shape_SurfaceArea (206) |  |  |
|  | log.sigma.3.0.mm.3D_glcm_DifferenceVariance (198) |  |  |
|  | original_shape_Flatness (166) |  |  |
|  | log.sigma.3.0.mm.3D_gldm_SmallDependenceLowGrayLevelEmphasis (154) |  |  |
|  | logarithm_firstorder_Maximum (148) |  |  |
|  | lbp.3D.m1_gldm_DependenceNonUniformity (139) |  |  |
| Ventral | original_shape_Maximum2DDiameterSlice (662) | 0.05 | 0.05 |
|  | original_shape_Maximum3DDiameter (633) |  |  |
|  | original_firstorder_Maximum (391) |  |  |
|  | log.sigma.1.0.mm.3D_gldm_SmallDependenceLowGrayLevelEmphasis (389) |  |  |
|  | log.sigma.3.0.mm.3D_firstorder_10Percentile (283) |  |  |
|  | log.sigma.5.0.mm.3D_glszm_SmallAreaHighGrayLevelEmphasis (249) |  |  |
|  | lbp.3D.m1_glszm_GrayLevelNonUniformity (211) |  |  |
|  | log.sigma.3.0.mm.3D_gldm_SmallDependenceLowGrayLevelEmphasis (177) |  |  |
|  | log.sigma.5.0.mm.3D_glszm_SmallAreaLowGrayLevelEmphasis (170) |  |  |
|  | log.sigma.5.0.mm.3D_glszm_ZonePercentage (166) |  |  |
| Lateral | original_shape_Maximum3DDiameter (563) | 0.19 | 0.001 |
|  | original_shape_Maximum2DDiameterSlice (514) |  |  |
|  | lbp.3D.m1_glszm_GrayLevelNonUniformityNormalized (422) |  |  |
|  | square_glszm_LargeAreaEmphasis (342) |  |  |
|  | lbp.3D.m1_firstorder_Range (313) |  |  |
|  | logarithm_firstorder_10Percentile (288) |  |  |
|  | log.sigma.3.0.mm.3D_gldm_SmallDependenceLowGrayLevelEmphasis (262) |  |  |
|  | original_shape_Sphericity (218) |  |  |
|  | lbp.3D.m1_gldm_DependenceEntropy (201) |  |  |
|  | logarithm_glcm_InverseVariance (189) |  |  |
| Intralaminar | original_shape_Maximum2DDiameterColumn (997) | 0.07 | 0.01 |
|  | lbp.3D.m1_glszm_GrayLevelNonUniformity (311) |  |  |
|  | original_shape_Maximum2DDiameterSlice (298) |  |  |
|  | lbp.3D.m2_firstorder_Minimum (260) |  |  |
|  | logarithm_firstorder_Mean (246) |  |  |
|  | log.sigma.3.0.mm.3D_glcm_ClusterTendency (230) |  |  |
|  | original_shape_MajorAxisLength (219) |  |  |
|  | lbp.3D.m1_gldm_SmallDependenceHighGrayLevelEmphasis (217) |  |  |
|  | lbp.3D.k_glszm_ZoneEntropy (196) |  |  |
|  | log.sigma.3.0.mm.3D_glcm_SumSquares (193) |  |  |

**References**

1- Tibshirani R. Regression Shrinkage and Selection via the Lasso. *J R Stat Soc*. 1996;58(1):267-288.
